# Liposomal Dexamethasone Reduces A/H1N1 Influenza-Associated Morbidity in Mice

**DOI:** 10.1101/2021.08.02.454850

**Authors:** Jung Won Kwon, Hailian Quan, Juha Song, Hyewon Chung, Daun Jung, Jung Joo Hong, Yi Rang Na, Seung Hyeok Seok

## Abstract

Re-emerging viral threats have continued to challenge the medical and public health systems. It has become clear that a significant number of severe viral infection cases are due to an overreaction of the immune system, which leads to hyperinflammation. In this study, we aimed to demonstrate the therapeutic efficacy of the dexamethasone nanomedicine in controlling the symptoms of influenza infection. We found that the A/Wisconsin/WSLH34939/2009 (H1N1) infection induced severe pneumonia in mice with a death rate of 80%, accompanied by significant epithelial cell damage, infiltration of immune cells, and accumulation of pro-inflammatory cytokines in the airway space. Moreover, the intranasal delivery of liposomal dexamethasone during disease progression reduced the death rate by 20%. It also significantly reduced the protein level of tumor necrosis factor-alpha (TNFα), interleukin1β (IL-1β), IL-6, and the C-X-C motif chemokine ligand 2 (CXCL2) as well as the number of infiltrated immune cells in the bronchoalveolar lavage fluids as compared to the free drug. It was found that the liposomal dexamethasone was mainly distributed into the monocyte/macrophages in the lungs, suggesting its mode of action via the specific delivery of the drug into myeloid cells as a major cell population for inducing the cytokine storm. Taken together, the intranasal delivery of liposomal dexamethasone may serve as a novel promising therapeutic strategy for the treatment of influenza A-induced pneumonia.

**Importance:** Influenza A virus causes annual epidemics and sporadic pandemics of respiratory diseases. An excessive immune response, called a cytokine storm, caused by the influenza virus is the most common complication of influenza infection that is associated with high levels of morbidity and mortality. Here, we report that the liposomal dexamethasone targeting the monocytes/macrophages during disease progression reduced the death rate as well as the protein level of inflammatory cytokines and the number of immune cells. Therefore, the findings of this study may be applied to address the limitations of the current strategies for the treatment of influenza.

## Introduction

Lower respiratory tract infections account for approximately 7% of infection-related deaths per year worldwide, and viruses are a common cause of community-acquired pneumonia (1–4). Among this wide group of viruses, influenza virus is of utmost importance, and numerous interventions have been proposed for its management, especially after the pandemic that was caused by the H1N1 influenza virus outbreak (5). The 2009 H1N1 influenza virus caused more than 18,000 confirmed deaths due to lung damage in the United States (6, 7). Almost 32% of patients either developed acute respiratory distress syndrome (ARDS) or died. These infections are accompanied by aggressive pro-inflammatory responses along with insufficient control of the anti-inflammatory responses, a combination of events that is referred to as the “cytokine storm (8, 9)”.

Following the primary exposure of respiratory epithelial cells to the influenza virus, progeny viruses that proliferate within these cells can infect other cells, including alveolar macrophages (10–12). The inflammatory response begins when the pathogen-associated molecular pattern (PAMP) of the virus is recognized by pattern recognition receptors (PRRs) of innate immune cells (13–15). Specific pro-inflammatory cytokines are expressed and lead to the recruitment of neutrophils, monocytes, macrophages, and T cells into the site of infection (11, 12). Hyperinflammation and macrophage activation syndrome (MAS) result in the overproduction of proinflammatory cytokines, such as the IL-1β, IL-6, and TNFα, as well as coagulation abnormalities, which contribute to organ failure and other fatalities (16–18). Therefore, the reduction of pro-inflammatory cytokine levels is as important as anti-viral therapies. Since monocytes and macrophages are the main cells that secrete pro-inflammatory cytokines, the efficient control of these cells can be used as a therapeutic target to alleviate inflammation.

Corticosteroids are a class of steroid hormones that exhibit anti-inflammatory activity by binding to the cytoplasmic corticosteroid receptor, which regulates the transcription of anti-inflammatory genes (19, 20). As the outcome of severe influenza is determined by both viral virulence and host resistance, the use of immunomodulatory therapy in combination with conventional antiviral therapy is highly warranted. During the 2009 H1N1 influenza pandemic, nearly 40% of patients in France were treated for ARDS using adjuvant systemic corticosteroids (21); however, the evidence supporting the use of corticosteroids in severe influenza was inconclusive. In this regard, Lammers et al. recently proposed the nano-formulation of dexamethasone within liposomes to improve the management of respiratory viral infections (22). Liposomes, microscopic phospholipid bubbles with a bilayered membrane structure, have been successfully applied in clinics as drug carriers to improve the delivery of drugs to target cells and tissues that play a key role in the acute and progressive phases of many diseases (23). Although dexamethasone nanomedicine has been used as a therapeutic agent for the coronavirus disease 2019 (COVID-19) (22), it has not yet been tested for the influenza infection in preclinical studies.

In this study, we demonstrated the therapeutic efficacy of liposomal dexamethasone (DEX/lipo) in a mouse model of influenza infection. We found that DEX/lipo outperformed the free drug in reducing the morbidity and mortality caused by the A/H1N1 influenza infection. Therefore, our findings offer valuable insights into the efficiency of the currently available corticosteroids for the treatment of influenza infection via the targeted delivery of these drugs.

## Results

### DEX/lipo is specifically delivered into the macrophages in the lungs

As monocytes and macrophages seem to be implicated in cytokine storm (24, 25), we encapsulated dexamethasone into liposomes with a particle size of 1,000 nm (DEX/lipo) (**Fig. 1A**) based on many other studies showing that liposomes with relatively larger diameters were more effectively distributed into phagocytes (26–28). After 24 hours of intranasal injection in mice, DiI-labeled DEX/lipo was detected in the cytosol of F4/80^+^ macrophages in the lungs (**Fig. 1B**). We further confirmed the distribution of injected liposomes as up to 70% of DiI^+^ cells were alveolar macrophages, interstitial macrophages and monocytes (**Fig. 1C**). In contrast, less than 10% of non-leukocytes and lymphocytes can uptake DiI^+^ liposomes. In particular, up to 15.8% of monocytes, 72.5% of alveolar macrophages, and 42.2% of interstitial macrophages in the lungs were DiI^+^ (**Fig. 1D**; gating schemes are shown in **Fig. S1**). Collectively, these results confirm that locally delivered DEX/lipo is mostly distributed into monocytes/macrophages in the lungs.

**Fig. 1.**
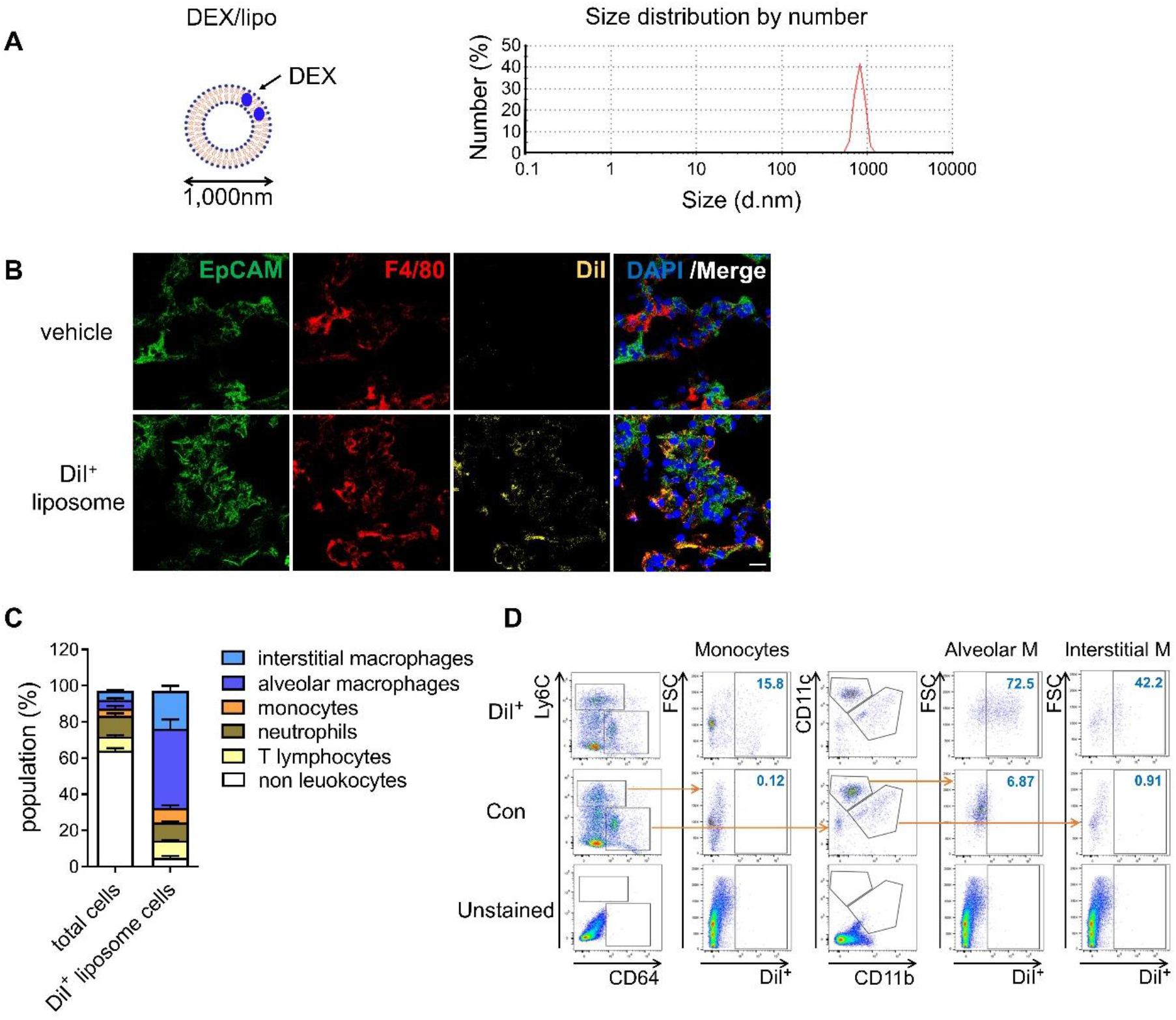
DEX/lipo is specifically delivered into the macrophages in lungs. (A) Schematic of DEX/lipo, left, and size determination of DEX/lipo by dynamic light scattering (DLS), right. The mean diameter is close to 1,000 nm. (B) Co-immunofluorescence (Co-IF) staining was performed to confirm the co-localization of the epithelial cell adhesion molecule (EpCAM) (green) or F4/80 (red) with DiI (yellow). Mice were injected intranally with 20 μl of DiI-labeled liposomes, and lung tissues were dissected at 24 h postinjection. Scale bar; 20 μm. (C) Population percentages in the lungs of mice are shown as total cells compared to DiI^+^ cells. Representatives of two independent experiments are shown. (D) Flow-cytometry analysis of DiI^+^ cells. CD45^+^Ly6C^+^CD64^low^ monocytes population, CD45^+^Ly6C^−^CD64^+^CD11c^+^ alveolar macrophages population, and CD45^+^Ly6C^−^CD64^+^CD11b^+^ interstitial macrophages population are shown in the DiI/forward scatter (FSC) dot plot with gating for the DiI^+^ population.

### DEX/lipo effectively reduces inflammation in mice infected with the lethal influenza A virus

We determined that DEX/lipo has a therapeutic effect on a mouse influenza model infected with influenza A/Wisconsin/WSLH34939/09. Mice under isoflurane anesthesia were infected intranally with 10^5^ tissue culture infectious dose 50 (TCID_50_) of influenza virus. One hour after infection, mice were anesthetized by isoflurane inhalation for intranasal delivery of PBS or dexamethasone (DEX) or DEX/lipo for three days. On day 3 postinfection, mice were treated with 100 μl of PBS, DEX, or DEX/lipo intravenously because of nasal watery. Mice were monitored for survival for 10 days (**Fig. 2A**). While 20% of PBS-treated mice survived on day 5 postinfection, mice treated with DEX/lipo showed the highest survival rate (40%). However, all mice treated with DEX died on day 5 (**Fig. 2B**). To verify the anti-inflammatory activity of DEX/lipo on the infiltration of inflammatory cells, mice were euthanized at day 10 postinfection to obtain lung tissues for histopathological examination. In mice treated with PBS, there was a severe inflammatory response characterized by alveolar congestion, thickening of the alveolar wall, and infiltration and aggregation of immune cells in airspaces or vessel walls. However, fewer inflammatory infiltrates were observed in the lungs treated with DEX/lipo than in the lungs of mice treated with PBS or DEX (**Fig. 2C**). Consistent with these findings, the total histopathological scores also decreased significantly in the DEX/lipo group (**Fig. 2D**). We also measured the virus titer to determine whether this histopathological score improvement in the DEX/lipo group was due to the difference in viral titers between groups. The viruses isolated from lungs plaqued on Madin-Darby canine kidney (MDCK) cells using TCID_50_. However, virus isolated from infected mouse lungs did not decrease in the DEX/lipo group (**Fig. 2E**). Collectively, these results support our hypothesis that targeting macrophages using DEX/lipo has a therapeutic effect in reducing inflammation.

**Fig. 2.**
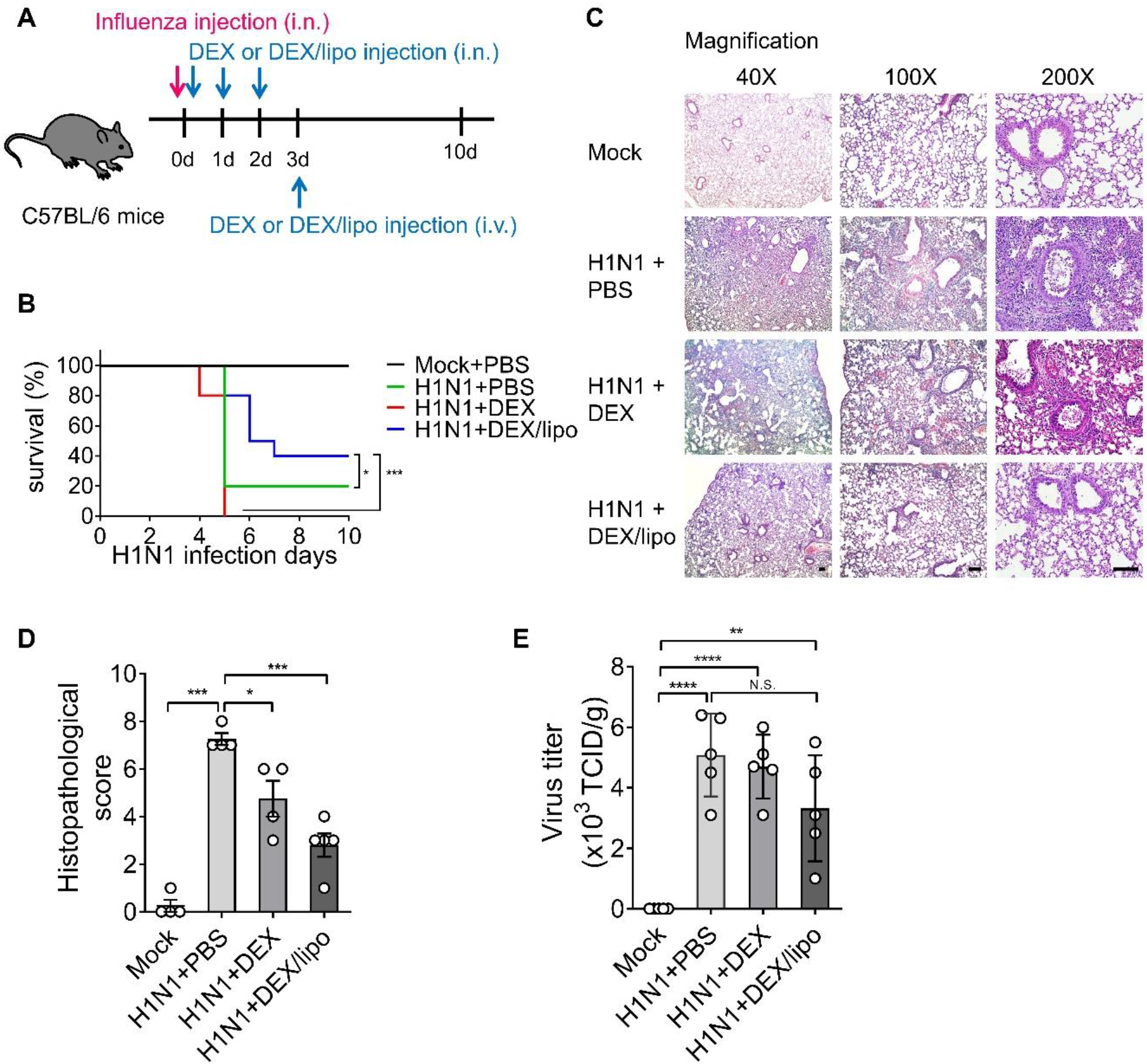
DEX/lipo effectively reduces inflammation in mice infected with the lethal influenza A virus. (A) Therapeutic regimen. Mice (n = 10 per group) were infected intranasally with 10^5^ TCID_50_ of influenza A virus (red arrow). One hour after infection, the mice were anesthetized by isoflurane inhalation for the intranasal delivery of the vehicle (phosphate-buffered saline (PBS)) or DEX (30 μg/kg) or DEX/lipo (30 μg/kg) for 3 days (blue arrow). On day 3 postinfection, the mice were injected intravenously with 100 μl of PBS or DEX or DEX/lipo due to nasal watering (blue arrow). (B) Survival rate of infected mice (n = 10 per group) after DEX or DEX/lipo treatment. \**P* < 0.05, \*\*\**P* < 0.001 by the log-rank test. (C) Hematoxylin and eosin staining of the lungs on day 10 postinfection. (n = 5 per group) Scale bar; 100 μm. (D) Comparison of the total histopathological scores of lung injury. (n = 4–5 per group) Data are presented as the mean ± standard error of the mean (SEM). \**P* < 0.05, \*\*\**P* < 0.001 by the Student’s t-test. (E) Lung viral titers from mice on day 10 postinfection using TCID_50_. (n = 4–5 per group) Data are presented as the mean ± SEM. \*\**P* < 0.01, \*\*\*\**P* < 0.0001 by the Student’s t-test.

### DEX/lipo decreases infiltrated cells in the bronchoalveolar lavage fluid of mice with lethal influenza infection

We analyzed the infiltrated cells in bronchoalveolar lavage fluid (BALF) collected from the lungs of mice infected with influenza A virus. Many inflammatory cells, such as monocytes, neutrophils, and lymphocytes, are recruited in the lungs of mice infected with influenza A virus. The cell infiltrates in the BALF of mice treated with PBS, DEX, or DEX/lipo were statistically analyzed for cell number and type (**Fig. 3**). DEX/lipo treatment greatly reduced the infiltration of macrophages, which contributed to acute lung injury in influenza pneumonia (29), and the infiltration of neutrophils and lymphocytes was reduced. Rather, DEX treatment increased the total number of infiltrated cells, especially macrophages and T cells, as compared to PBS treatment (**Fig. 3B-D**).

**Fig. 3.**
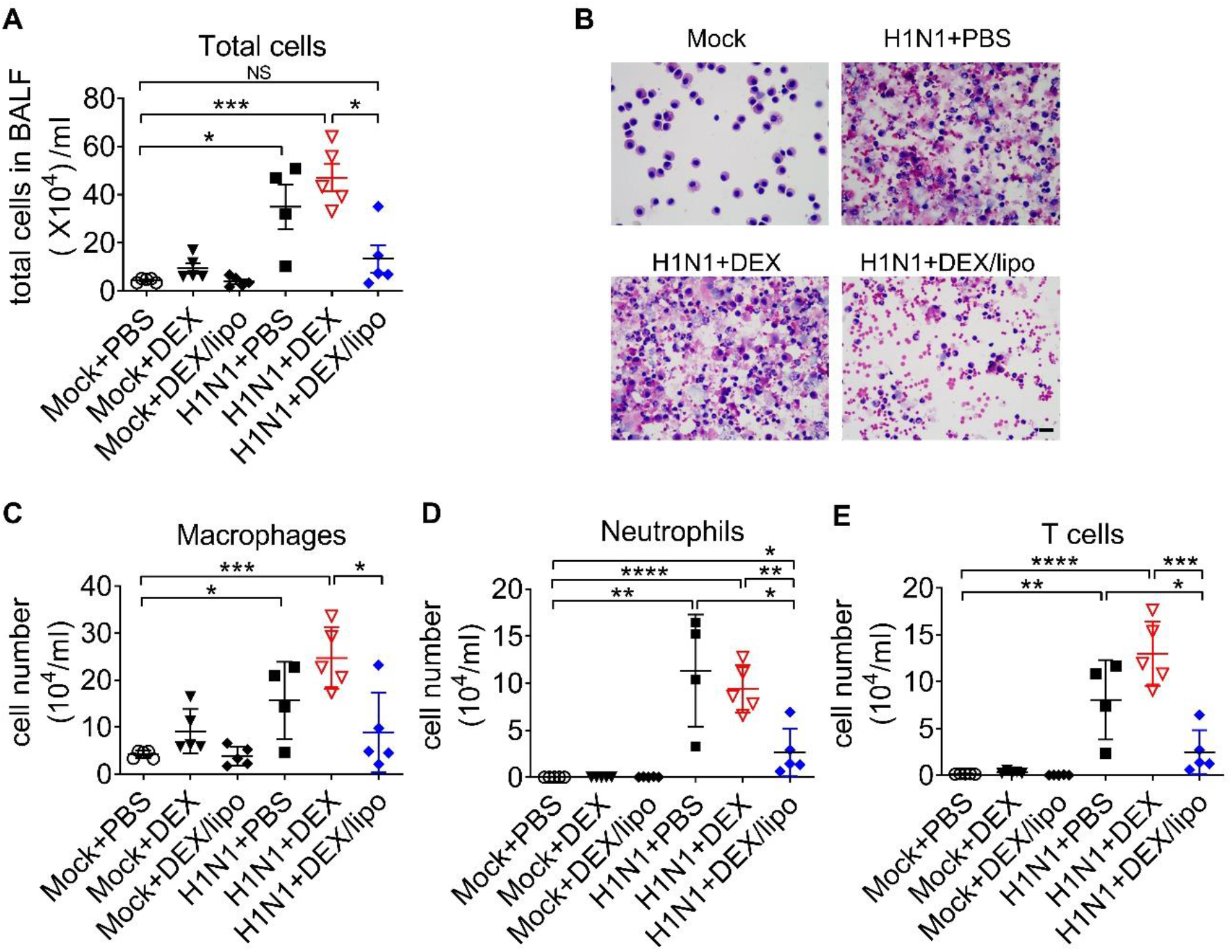
DEX/lipo decreases the total number of cells in the bronchoalveolar lavage fluid of mice with lethal influenza infection. (A) Analysis of the total number of infiltrated cells in the BALF collected from the mock or influenza A-virus infected mice treated with PBS or DEX or DEX/lipo (n = 4–5 per group) on day 2 postinfection. (B) Differential cell counts of BALF cells. Representative pictures of Diff-quick staining of cytospin preparation. Scale bar; 20 μm Changes in the number of macrophages (C), neutrophils (D), T cells (E). BALF cells counted using the QWin program (Leica Microsystems). Data are presented as the mean ± SEM and are representative of two independent experiments. \**P* < 0.05, \*\**P* < 0.01, \*\*\**P* < 0.001, \*\*\*\**P* < 0.0001 by the Student’s t-test.

### DEX/lipo significantly reduces the pro-inflammatory cytokines and chemokines in mice with lethal influenza infection

Given that inflammatory cytokines and chemokines are linked to lung damage in severe influenza pneumonia, we determined the role of DEX/lipo in mice with influenza infection. The protein levels of pro-inflammatory cytokines and chemokines, TNFα, IL-1β, IL-6, CXCL1, and CXCL2 increased significantly during influenza infection (**Fig. 4 A-E**). Consistent with the infiltrated total cell number data, cytokines and chemokines were increased in the H1N1+DEX group, except for CXCL1. In particular, IL-1β was significantly increased compared to that in the H1N1+PBS group. Production of TNFα, IL-1β, IL-6, and CXCL2 was reduced in the H1N1+DEX/lipo group, but CXCL1 was not affected compared to the H1N1+DEX group (**Fig. 4 A-E**). In addition, DEX/lipo treatment reduced TNFα, IL-1β, and IL-6 to a level similar to that of the mock treatment (**Fig. 4 A-C**).

**Fig. 4.**
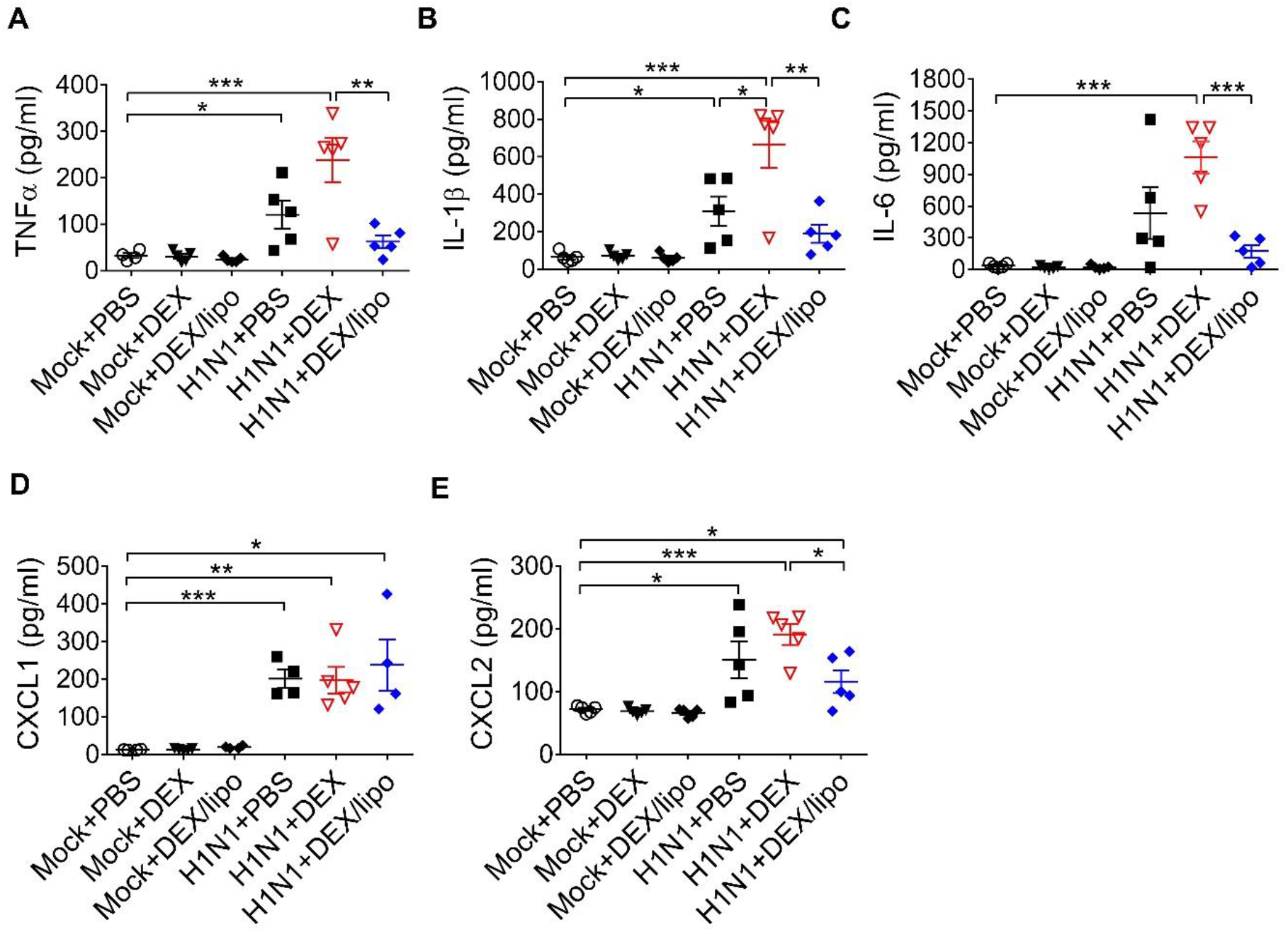
DEX/lipo significantly reduces the pro-inflammatory cytokines and chemokines in mice with lethal influenza infection. (A-E) Analysis of the inflammatory cytokines and chemokines collected from lungs of mock or influenza A virus infected mice treated with PBS or DEX or DEX/lipo (n = 4–5 per group) on day 2 postinfection. TNFα (A), IL-1β (B), IL-6 (C), CXCL-1 (D), and CXCL2 (E) were measured by the enzyme-linked immunosorbent assay (ELISA). Data are presented as the means ± SEM and are representative of two independent experiments. \**P* < 0.05, \*\**P* < 0.01, \*\*\**P* < 0.001 by the Student’s t-test.

## Discussion

Many patients with severe influenza die from overwhelming viral pneumonia and serious complications caused by cytokine storms. In this study, we revealed that DEX/lipo effectively reduces pro-inflammatory responses in hosts with pathogenic A/H1N1 influenza infection, demonstrating the possible availability of DEX/lipo in controlling disease symptoms of cytokine storm in respiratory infections.

Pathogenesis in human influenza infection is accompanied by diffuse epithelial sloughing in tracheal, bronchial, and bronchiole biopsy, predominant presence of mononuclear cells, and bloody exudate in the airway lumen with interstitial swelling (30, 31). In line with this, a mouse model of human pathogenic pandemic A/Wisconsin/*WSLH34939/09* influenza virus infection previously showed 80% of death within 12 days post-infection with marked tissue injury, including hemorrhage, pulmonary edema, and mononuclear cell accumulation (6). The mouse adapted influenza infection model established in this study showed a faster pathogenesis than the previous study, showing 80% death on day 5 postinfection, but the characteristic histopathological findings, including alveolar congestion and infiltration of macrophages in airway spaces, showed comparable results (**Fig. 2, Fig. 3**). Of note, cytokines, such as TNFα and IL-6, all of which are known to be directly correlated with host morbidity and pulmonary injury (32–34), were significantly elevated in bronchoalveolar lavage fluids on day 2 postinfection (**Fig. 4A and C**), implying the importance of immune responses in the disease pathogenesis of our influenza infection model.

The most notable results in this study were the differences in the therapeutic responses of DEX/lipo compared to the free drug. Intranasal delivery of free dexamethasone did not alleviate tissue injury in the infected mice (**Fig. 2C**). This was further supported by the results shown in Fig. 3 and 4, because free dexamethasone did not block the accumulation of macrophages and T cells or decrease of TNFα, IL-1β, IL-6, and CXCL2 in bronchoalveolar fluids. In fact, free dexamethasone accelerated the fatality rate of infected mice from 80% to 100% (**Fig. 2B**), possibly due to increased T cells and IL-1β in airway spaces. In contrast, DEX/lipo treatment rescued infected mice from death compared to free drug, with a significant decrease in tissue damage, infiltration of macrophages and T cells, and accumulation of TNFα, IL-1β, IL-6, and CXCL2 in the lungs. These results are consistent with concerns from previous observations of dexamethasone toxicities in the clinic (35–37), and further demonstrate the usability of liposomes as delivery cargo in improving the safety and effectiveness of dexamethasone in controlling influenza infection.

There have been many cases in which DEX/lipo, mainly targeting macrophages, was delivered to reduce inflammation in various diseases. Polyethylene glycol (PEG)-free formulation of macrophage-targeting DEX/lipo reduces the dose and/or frequency required to treat adjuvant arthritis, with the potential to enhance or prolong therapeutic efficacy and limit side effects in the treatment of rheumatoid arthritis (38). Additionally, administration of tumor-associated macrophage (TAM)-targeting DEX/lipo resulted in a significant inhibition of tumor growth and metastasis in a model of prostate cancer bone metastases (39). In particular, macrophages are a major producer of inflammatory cytokines in influenza infection, and we showed that DEX/lipo were effectively distributed into monocytes/macrophages (**Fig. 2C**). The fact that DEX/lipo did not reduce CXCL1, which is mostly produced by epithelial cells, and not by macrophages (**Fig. 4D**), further supports our hypothesis that liposomal encapsulation specifically delivered dexamethasone into macrophages, but not epithelial cells.

Antiviral therapy should be accompanied by an immunomodulatory agent because antiviral drugs do not directly inhibit inflammation in pulmonary injury. In our study, even with the single treatment of DEX/lipo, decreased inflammatory cytokines, and increased the survival rate.

In conclusion, our data demonstrate that the macrophage-targeting DEX/lipo plays crucial roles in the prevention of pneumonia induced by influenza as well as the reduction of pro-inflammatory cytokines/chemokines and infiltration of inflammatory cells (**Fig. 5**). Therefore, targeting macrophages using DEX/lipo may be used as a promising therapeutic approach for the treatment of cytokine storm-induced influenza infection with antiviral drugs.

**Fig. 5.**
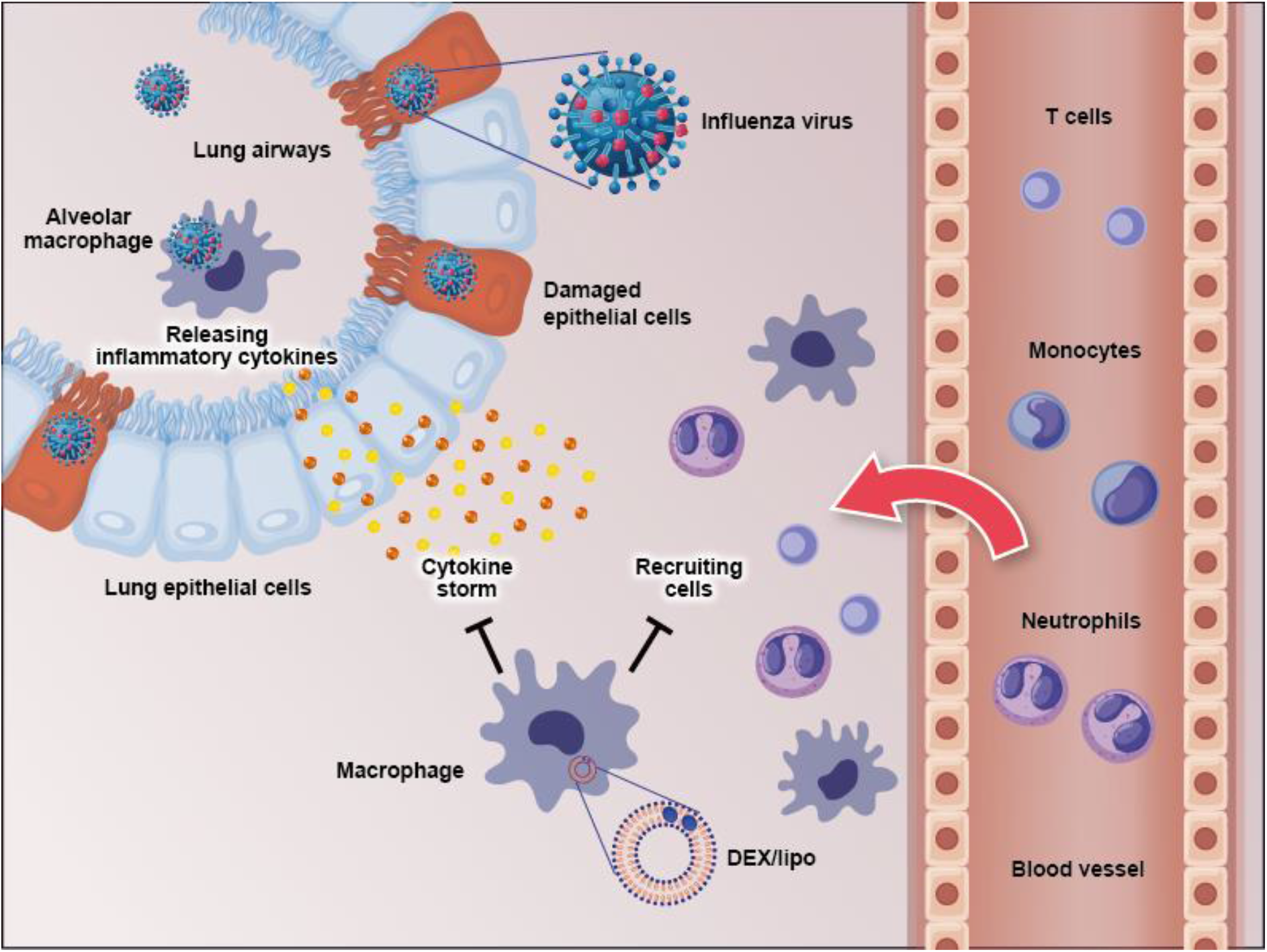
Schematic figure of the effects of DEX/lipo on the pathogenesis of the influenza virus infection. Influenza virus infects the lung epithelial cells and subsequently alveolar macrophages. This results in pro-inflammatory cytokine, and chemokine secretion by mainly macrophages. Specific pro-inflammatory cytokines lead to the recruitment of neutrophils, monocytes, macrophages, and T cells into the site of infection. DEX/lipo delivery targeting macrophages reduces cytokines/chemokines secretion and recruitment of leukocytes to the lung of mice with lethal influenza infection.

## Materials and Methods

### Mice, virus, and cell line

Male C57BL/6J mice of 7-weeks-of-age were purchased from ORIENT BIO (Orient Bio, Gyeonggi-do, Korea). Animal experiments were conducted at the Institute for Experimental Animals, College of Medicine and cared for according to the Guide for the Care and Use of Laboratory Animals prepared by the Institutional Animal Care and Use Committee of Seoul National University at Seoul (accession number SNU-160307-6-1). Mice were housed in cages with a constant-flow air exchange, supporting specific pathogen-free conditions. Influenza A/Wisconsin/*WSLH34939*/09 was obtained from the Michael laboratory at the Scripps Research Institute. The adaptation method for influenza A virus in mouse lungs is as follows. The C57BL/6J mice were each anesthetized with isoflurane (HANA PHARM CO., Gyeonggi-do, Korea), intranasally inoculated with 10^5^ TCID_50_ of influenza A/Wisconsin/*WSLH34939*/09 virus in a volume of 20 μl. At day 2 postinfection, the lungs from mice were homogenized in the Dulbecco’s modified Eagle medium (DMEM) medium with 10% fetal bovine serum (FBS) and 1% penicillin-streptomycin (PS), centrifuged at 9,520 × *g* for 10 min, and the supernatant was stored at −80 °C. The first adapted virus was amplified and plaqued on the MDCK cells using TCID_50_. Briefly, the assay was performed by adding a serial dilution of the virus sample to cells in a 96-well plate. After incubation, the percentage of infected wells was observed for each dilution, and the results were used to calculate the TCID_50_ value. This calculation can generally be performed using the Spearman-Karber method. MDCK cells were cultured in DMEM supplemented with 10% FBS and 1% PS.

Mice under isoflurane anesthesia were infected intranasally with 10^5^ TCID_50_ of the adapted influenza virus A. One hour after infection, the mice were anesthetized by isoflurane inhalation for the intranasal delivery of vehicle (20 μl of PBS) or dexamethasone (30 μg/kg dissolved in PBS) or DEX/lipo (30 μg/kg encapsulated in liposomes) for 3 days. On day 3 postinfection, the mice were injected intravenously with 100 μl of PBS, dexamethasone, or DEX/lipo due to nasal watering. The mice monitored for survival were euthanized when 25% of their starting weight was lost.

### Reagents

Dexamethasone were purchased from Sigma (Sigma, St. Louis, MO).

### Preparation of liposomes

Dexamethasone was encapsulated into liposomes with a diameter of 1,000 nm at a concentration of 1 mg/ml. Briefly, L-alpha-phosphatidylcholine (Sigma, St. Louis, MO) and cholesterol (Sigma, St. Louis, MO) were dissolved in a 2:1 mixture of chloroform and methanol and dried under a nitrogen stream, followed by a vacuum pump. The lipid film was hydrolyzed with PBS buffer and sized to 1,000 nm in a mini-extruder equipped with a 1,000 nm pore polycarbonate membrane (Avanti Polar Lipids, Alabama, AL). The diameter of liposomes was checked by dynamic light scattering (DLS). Liposomes were labeled with DiI (Molecular Probes, Eugene, OR) for visual detection.

### Flow cytometry

The mice lungs were obtained at 24 h postinjection with DiI-stained liposomes and digested with collagenase type I (Sigma, St. Louis, MO) to obtain single cells. Single cells were prepared for flow cytometric analysis. Anti-mouse CD16/32 antibody (clone number 93) was pre-added to block the non-specific binding of immunoglobulin to macrophage Fc receptors.

For surface marker analysis, live cells were re-suspended in staining buffer (1% bovine serum albumin (BSA), 5 mM ethylenediaminetetraacetic acid (EDTA), and 0.1% NaN_3_ in PBS) and stained with anti-mouse CD45 (30-F11), F4/80 (BM8), CD11b (M1/70), CD3 (145-2C11), CD64 (X54-5/7.1; BD Biosciences), Ly6C (HK1.4), and Ly6G (1A8-Ly6g) antibodies at 4 °C for 20 min. All antibodies were obtained from eBioscience, unless otherwise indicated. Data were acquired using LSR Fortessa (BD Biosciences) and analyzed using the FlowJo software (FlowJo LCC, Ashland, OR).

### Immunofluorescence (IF) analysis

The mice lungs were obtained at 24 h postinjection with DiI-stained liposomes. Frozen tissues were sectioned to a thickness of 5 μm and fixed with 4% paraformaldehyde (Merck, Kenilworth, NJ). Slides were incubated with 1% BSA in PBST for 1 h to block the non-specific antibody binding. Primary antibodies were pre-diluted in blocking buffer to 1:200 for F4/80 (CI-A3-1; Abcam) or EpCAM (G8.8; eBioscience) and applied to tissue sections overnight at 4 °C in a humidified chamber. The next day, appropriate secondary antibodies were applied, and nuclei were stained with 4,6-diamidino-2-phenylindole (DAPI; Invitrogen, Waltham, MA) before mounting. Fluorescence signals were detected using a Leica TCS SP8 confocal microscope (Leica, Buffalo Grove, IL).

### Cytokine and chemokine analyses

On day 2 postinfection, the trachea of euthanized mice was exposed, transected, and intubated with a blunt 18-gage needle that delivered 0.8 ml ice-cold PBS. Infusion of the 0.5 ml volume was repeated twice, and the fluid was recovered. The recovered BALF was centrifuged at 3,000 × *g* for 3 min at 4 °C and stored at −80 °C until further use. ELISAs were performed on the BALF using TNFα, IL-6 ELISA kits (BD Biosciences, Palo Alto, CA), IL-1β, CXCL1, and CXCL2 Duoset ELISA kits (R&D systems, Minneapolis, MN).

### Cell counting

BALF was obtained via cannulation of the trachea and lavaging the airway lumen with 0.8 ml ice-cold PBS for three times on day 2 postinfection. The recovered fluid was centrifuged, and the cell pellets were resuspended in PBS to count the total number of cells. Differential cell counts in the BALF were performed using the Diff-Quik staining reagent (Sysmex, Osaki, Tokyo), according to the manufacturer’s instructions. The numbers of macrophages, neutrophils, and T cells were calculated by multiplying the percentages obtained by the total yield. Slides were imaged using the QWin program (Leica, Leider Lane Buffalo Grove, IL).

### Histopathological scoring

Lung tissue samples were fixed in 4% paraformaldehyde neutral buffer solution for 24 h, dehydrated in a graded ethanol series, embedded in paraffin, sliced at 5 μm, and stained with hematoxylin and eosin. Lung histopathological score was assessed using the following parameters: a scale of 0 to 3 (0 = absent and appeared normal, 1 = light, 2 = moderate, and 3 = severe) according to the histologic features: 1) edema, hyperemia, and congestion; 2) neutrophil margination and tissue infiltration; 3) intra-alveolar hemorrhage and debris; and 4) cellular hyperplasia. Each parameter was scored from 0–3 based on severity. The total score was used to assess lung injury, which was calculated as the sum of all scores for each parameter (0–3, normal to minimal injury; 4,6, mild injury; 7,9, moderate injury; and 10,12, severe injury). The maximum score per animal was 12. Lung section scoring was performed at a low power (× 40).

### Quantification and statistical analyses

GraphPad Prism v.8 (GraphPad Software, La Jolla, CA) was used for the statistical analysis. Results are presented as the mean ± SEM for the experiments, unless otherwise indicated. Unpaired two-tailed Student’s t-tests were used to compare the two groups of independent samples. Sample size (n) is indicated in the figure legends. Statistical significance was set at *P* values of less than 0.05.

## Acknowledgements

This study was supported by Basic Science Research Program through the National Research Foundation of Korea (NRF) funded by the Ministry of Science, ICT & Future Planning (2020R1A2C2010202), the Korea Research Institute of Bioscience and Biotechnology Research Initiative Program (KGM4572121), the NRF (NRF-2021R1A2C2010219), and SUNH Research Fund (0320210190).

---

**Fig. S1.**
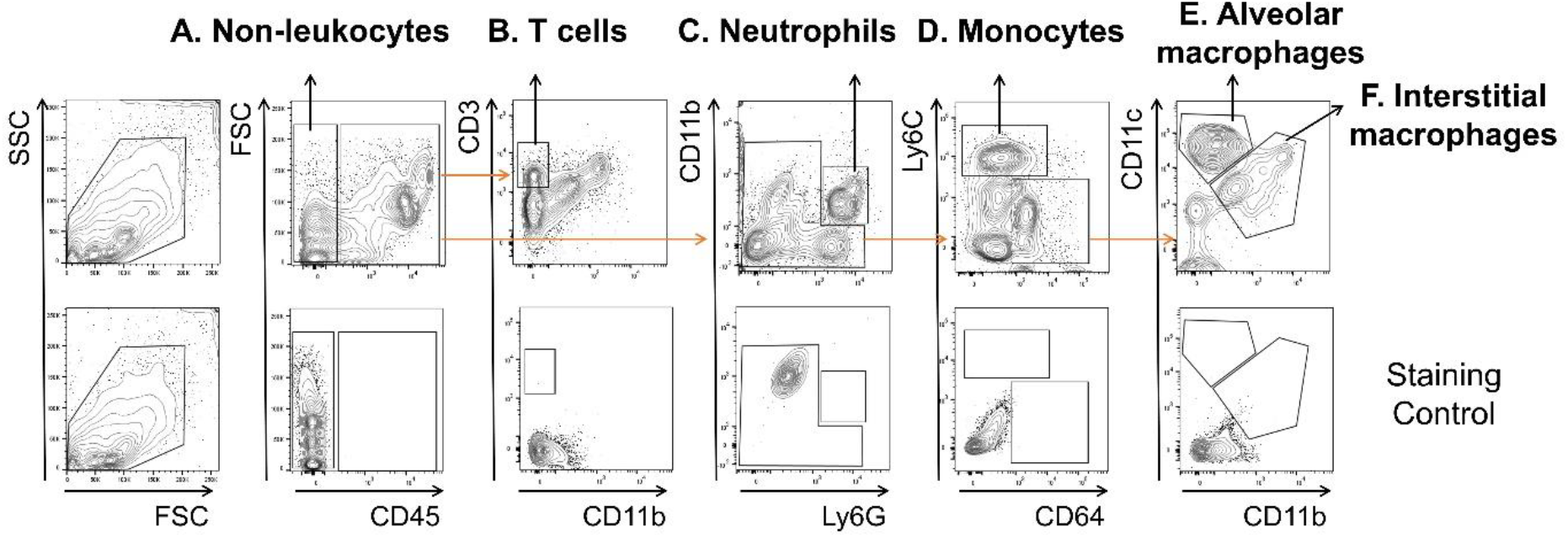
Gating strategies to determine the cellular compositions of the lungs of mice (Related to Fig. 1). Single cells were isolated from the lungs of DiI^+^ injected mice and analyzed by flow cytometry. Sequentially gated cells were (A) CD)-45^−^non-leukocytes, (B) CD45^+^ CD11b^−^CD3^+^ T cells, (C) CD45^+^ CD11b^+^Ly6G^+^ neutrophils, (D) CD45^+^ Ly6G^−^CD64^low^ Ly6C^+^ monocytes, (E) CD45^+^ Ly6G^−^CD64^+^ CD11c^+^ alveolar macrophages, (F) CD45^+^ Ly6G^−^CD64^+^ CD11b^+^ interstitial macrophages. Representative dot plots are shown.

